# Response patterns and mechanisms of plants to water stress

**DOI:** 10.1101/2020.03.02.973610

**Authors:** Yuan Sun, Cuiting Wang, Han Y.H. Chen, Honghua Ruan

## Abstract

Plants are key to the functionality of many ecosystem processes. The duration and intensity of water stress are anticipated to increase in the future; however, an elucidation of the responses of plants to water stress remains incomplete. For this study, we present a global meta-analysis derived from 1301 paired observations from 84 studies to evaluate the response patterns and mechanisms of plants to water stress. The results revealed that while water stress inhibited plant growth and photosynthesis, reactive oxygen species (ROS), plasma membrane permeability, enzymatic antioxidants, and non-enzymatic antioxidants increased. These responses generally increased with the intensity of water stress but were mitigated with experimental duration. Our findings suggested that the overproduction of ROS was the primary mechanism of plants in response to water stress and that plants tend to acclimate to water stress over time to some extent. Our synthesis provides a framework for understanding the responses and mechanisms of plants under drought conditions.

**One senence summary:** The overproduction of ROS was the primary mechanism of plants in response to water stress and that plants tend to acclimate to water stress over time to some extent.

## 1. Introduction

Drought is expected to increase continuously and significantly by the end of this century (Choat et al., 2012; IPCC, 2013). Water stress is problematic for plant growth and development (McDowell et al., 2011), as it limits access to resources required for photosynthesis due to stomatal closure and reduced internal water transport (Breda et al., 2006). As such, water stress impairs normal plant functionality and further induces morphological, physiological, and biochemical changes to compensate for water limitations (Mitchell et al., 2013; Lee et al., 2016). Understanding the patterns and mechanisms of responses by plants to water stress is central to predicting future plant functionality and resilience to drought episodes.

The impacts of water stress on plant growth, physiology, and biochemistry are well documented, and numerous individual studies have examined the roles of plant physiological indexes as relates to their tolerance to water stress (van der Molen et al., 2011; Zwicke et al., 2015). For plants, water limitations lead to the overproduction of reactive oxygen species (ROS), such as hydrogen peroxide (H_2_O_2_), and superoxide anion radical (O_2_^-.^), which results in growth inhibition (Wallace et al., 2016), decreases in photosynthetic functions (Deeba et al., 2012), lipid peroxidation, and further programmed cell death processes (Gill and Tuteja, 2010). However, to adapt to water stress, plants have evolved many acclimation mechanisms, including osmotic adjustment and antioxidant defense systems, which may enhance their ability to grow and develop under drought conditions (Fu and Huang, 2001; Khaleghi et al., 2019). Under water stress conditions, soluble sugars and proline accumulate to serve as osmolytes in various plants, assist in membrane protein stabilization, and ultimately increase plant resistance against water stress (Ashraf and Foolad, 2007; Gomes et al., 2010; Per et al., 2017). Further, ROS scavenging enzymatic antioxidants, such as superoxide dismutase (SOD), peroxidase (POD), catalase (CAT), glutathione reductase (GR), and ascorbate peroxidase (APX) can be activated to clear these excessive ROS (Gill and Tuteja, 2010). Adjustments in the activities of these enzymes are likely the primary path in plants for tolerating water stress (Nikoleta-Kleio et al., 2020).

The challenge remains to comprehensively address how various plants respond to water stress, as this can vary considerably (Rigui et al., 2019). For example, the ROS in leaves may increase (Tang et al., 2017) or decrease (Saglam et al., 2011) under water stress. Similarly, water stress can enhance (Sedaghat et al., 2017) or depress (Zhang et al., 2017) the SOD activity of plants. Previous studies have revealed that the performance of plant responses to water stress may decrease with experimental intensity and duration (Schneider et al., 2018), and vary with different plant species and tissues (Mirzaee et al., 2013; Lum et al., 2014). Therefore, it was necessary to conduct a systematic analysis to summarize the responses of plants under water stress.

The meta-analysis is a statistical methodology for synthesizing results across studies to reach an overall understanding of a problem (Gurevitch et al., 2018). A recent meta-analysis has specifically addressed the responses of plants to drought stress (Dong et al., 2017), but they focused on the physiological indexes (i.e., plant height, proline, electrolyte leakage, and root length) associated with transcription factors C-repeat/dehydration-responsive element-binding proteins which play important roles in plant response to environmental perturbations. Here we focus on the responses of ROS and enzymatic antioxidants (SOD, POD, CAT, GR, and APX), which represent defense mechanisms for plants under abiotic stresses (Gill and Tuteja, 2010; Sun et al., 2019). For this study, we established a global dataset by retrieving published papers to January 2020, including 1301 water-stress experiments from 84 papers (Table S1). Our objectives were to explore the general patterns and mechanisms of plants to water stress with the aim of providing reliable physiological indexes for the screening of drought-resistant species in the future.

## 2. Materials and Methods

### 2.1 Data collection

The database utilized in this meta-analysis was collected from peer-reviewed publications (Table S1) via the Web of Science and Google Scholar, prior to February 2020. The publication screening process is provided in Fig. S1. Our search terms were “water stress” or “water reduction”, or “drought” and “plant”. The following criteria were applied in this investigation: (1) Water stress and control groups began under the same abiotic and biotic conditions. (2) If the experiment included additional treatments, data were selected from the control and water stress groups only. (3) The water stress manipulation technique and duration had to be reported. (4) The sample sizes and means for the control and treatment groups were directly reported or could be extracted using WebPlotDigitizer (Burda et al., 2017). Our final dataset included 1301 paired observations from 84 primary articles (Fig. 1).

**Figure 1.**
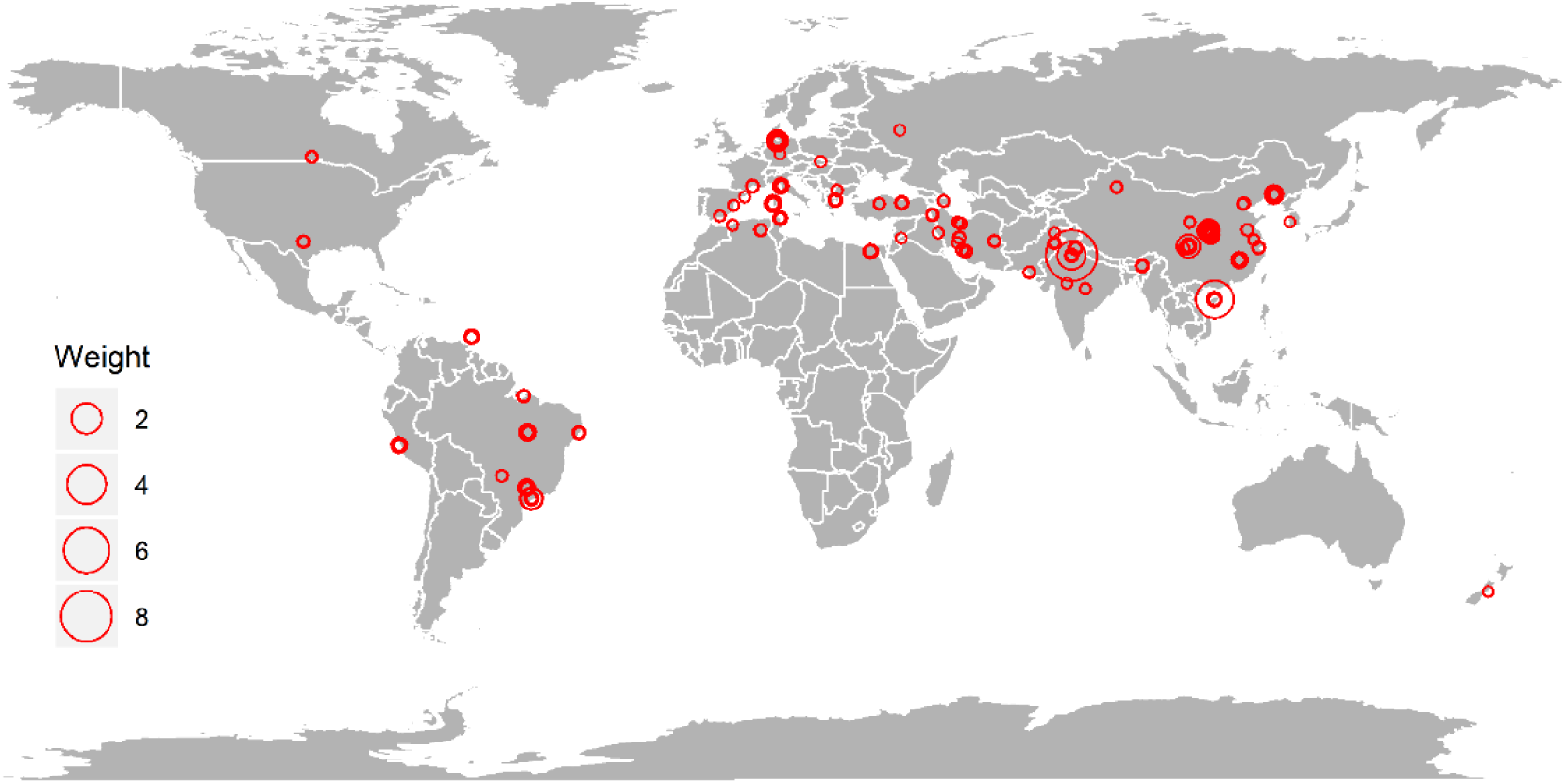
Geographic distribution of sites, with dot sizes representing study weight.

Eighteen physiological indexes, including abscisic acid (ABA), ascorbate peroxidase (APX), ascorbate (AsA), carotenoid (Car), CAT, chlorophyll (Chl), dry weight, electrolyte leakage (EL), maximal efficiency of PSII photochemistry (Fv/Fm), glutathione reductase (GR), malondialdehyde (MDA), POD, proline, protein, photochemical quenching coefficient (qP), ROS, SOD, and soluble sugar were collected. Specifically, we combined different indexes into specific plant performance, i.e., dry weight and protein into growth; Chl, Fv/Fm, and qP into photosynthesis (PS); ROS, MDA, and EL into plasma membrane permeability (PMP); APX, CAT, POD, SOD, and GR into enzymatic antioxidants (EA); and proline, soluble sugar, ABA, AsA, and Car into non-enzymatic antioxidants (NEA) based on morphology, physiology, and functionalities of plants (Gill and Tuteja, 2010). Several independent variables that might affect these responses variables were also collected, encompassing plant tissues tested indexes (whole plant, leaf, shoots, and roots), a mean water stress intensity of 0.52 (0.05-0.88), and a mean experimental duration of 36 days (1-365 days). Water stress intensity was calculated as the proportional reduction in soil moisture (reduced soil moisture under water stress treatment/soil moisture in the control groups) and experimental duration was the number of days since the experiment.

### 2.2 Statistical analyses

We employed natural log response ratios (lnRR) as effect sizes (Hedges et al., 1999) to estimate the magnitude of the treatment effect. The lnRR was calculated as ln (X_i_/X_c_) = lnX_i_ - lnX_c_, where X_i_ and X_c_ are the mean values for the water stress and control groups, respectively. The lnRR was weighted by the reciprocal of sampling variance, which was calculated as ln [(1/n_i_) × (S_i_/X_i_)^2^ + (1/n_c_) × (S_c_/X_c_)^2^] using the R package *metafor* 2.1.0 (Viechtbauer, 2010), where S_i_ and S_c_ represent the standard deviations of the water stress and control groups, respectively, with n_i_ and n_c_ as sample sizes. In instances where the standard deviations (SD) were not reported (a total of 86 observations) we imputed them using the “Bracken 1992” method (Benitez-Lopez et al., 2017; Sun et al., 2019) with *metagear* (Lajeunesse, 2016).

For each plant parameter, we used the following linear mixed-effect model to test whether the mean lnRR differed from zero:

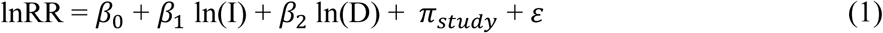

where I and D represent the water stress intensity and experimental duration. *β*_*n*_, *π*_*study*_, and ε are the coefficients to be calculated, the random effect factor of “Study”, and sampling error, respectively. We applied linear mixed-effects models using the restricted maximum likelihood estimation with the *lme4* package (Bates et al., 2015). Continuous predictors including water stress intensity and experimental duration in Equation (1) were scaled (observed minus mean and divided by one SD).

To examine the linearity assumption between dependent and independent variables, we compared the logarithmic and linear functions for I and D using the *MuMIn* package (Bartoń, 2018), and found that the logarithmic functions for I and D resulted in lower, or similar, Akaike information criterion (AIC) values (Table S2). For consistency, we analyzed variables with Equation (1).

To maximize the comparability, we tested the effects of plant performances and tissues on lnRR by adding plant performances, and tissues to Equation (1). For ease of interpretation, we transformed the lnRR and its corresponding confidence interval (CI) using [exp (lnRR)-1] × 100%. Further, linear-regressions were employed to examine the correlations of index ratio responses with water stress intensity and experimental duration. All statistical analyses were performed using R 3.6.0 software (R Development Core Team, 2019).

## 3. Results

Across all individual studies, the ABA increased significantly, by 126.6% on average (CI, 26.9-226.3%; *P* = 0.01), AsA by 19.3% (9.1-29.5%; *P* < 0.01), CAT by 28.8% (14.3-43.4%, *P* < 0.01), EL by 99.4% (45.9-153.0%, *P* < 0.01), MDA by 44.2% (19.9-68.5%, *P* < 0.01), POD by 28.0% (11.7-44.2%, *P* < 0.01), proline by 136.8% (59.9-213.7%, *P* < 0.001), ROS by 65.7% (33.8-97.6%, *P* < 0.001), SOD by 29.8% (15.4-44.1%, *P* < 0.001), and soluble sugar by 116.9% (32.2-201.5%, *P* = 0.03) under water stress, compared to the mean of the control groups (Fig. 2). However, on average, water stress significantly (*P* < 0.05) decreased Chl by 32.5%, dry weight by 37.4%, Fv/Fm by 13.1%, and qP by 50.2%, but had no significant impacts on APX, Car, GR, and protein (all *P* > 0.05; Fig. 2).

**Figure 2.**
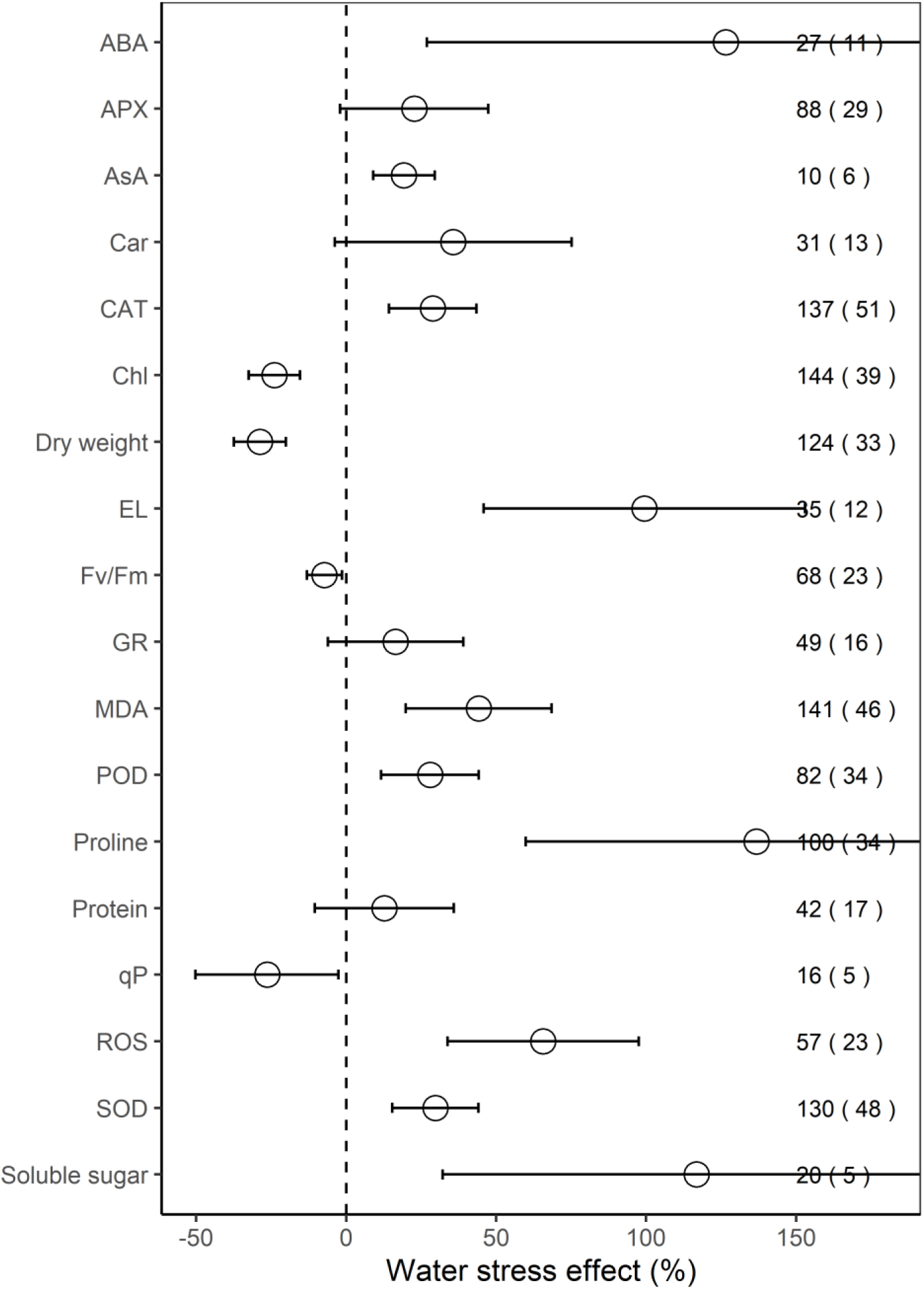
The response of physiological indexes to water stress. Values are the means and 95% confidence intervals. The dashed black line is zero effect size. Numbers without and within parentheses represent the number of observations and studies, respectively. ABA, APX, AsA, Car, CAT, Chl, EL, Fv/Fm, GR, MDA, POD, qP, ROS, and SOD represent abscisic acid, ascorbate peroxidase, ascorbate, carotenoid, catalase, chlorophyll, electrolyte leakage, maximal efficiency of PSII photochemistry, glutathione reductase, malondialdehyde, peroxidase, photochemical quenching coefficient, reactive oxygen species, and superoxide dismutase, respectively.

We found that the effect sizes for EA, NEA, and PMP increased significantly under water stress (all *P* < 0.001), and the effect size for growth and PS decreased (all *P* < 0.001; Fig. 3a). Furthermore, for plant tissues tested indexes, water stress had positive effects on leaves (16.9%, *P* < 0.01), negative effects on shoots (−37.5%, *P* < 0.001), but no effects on the whole plants and roots (all *P* > 0.05; Fig. 3b).

**Figure 3.**
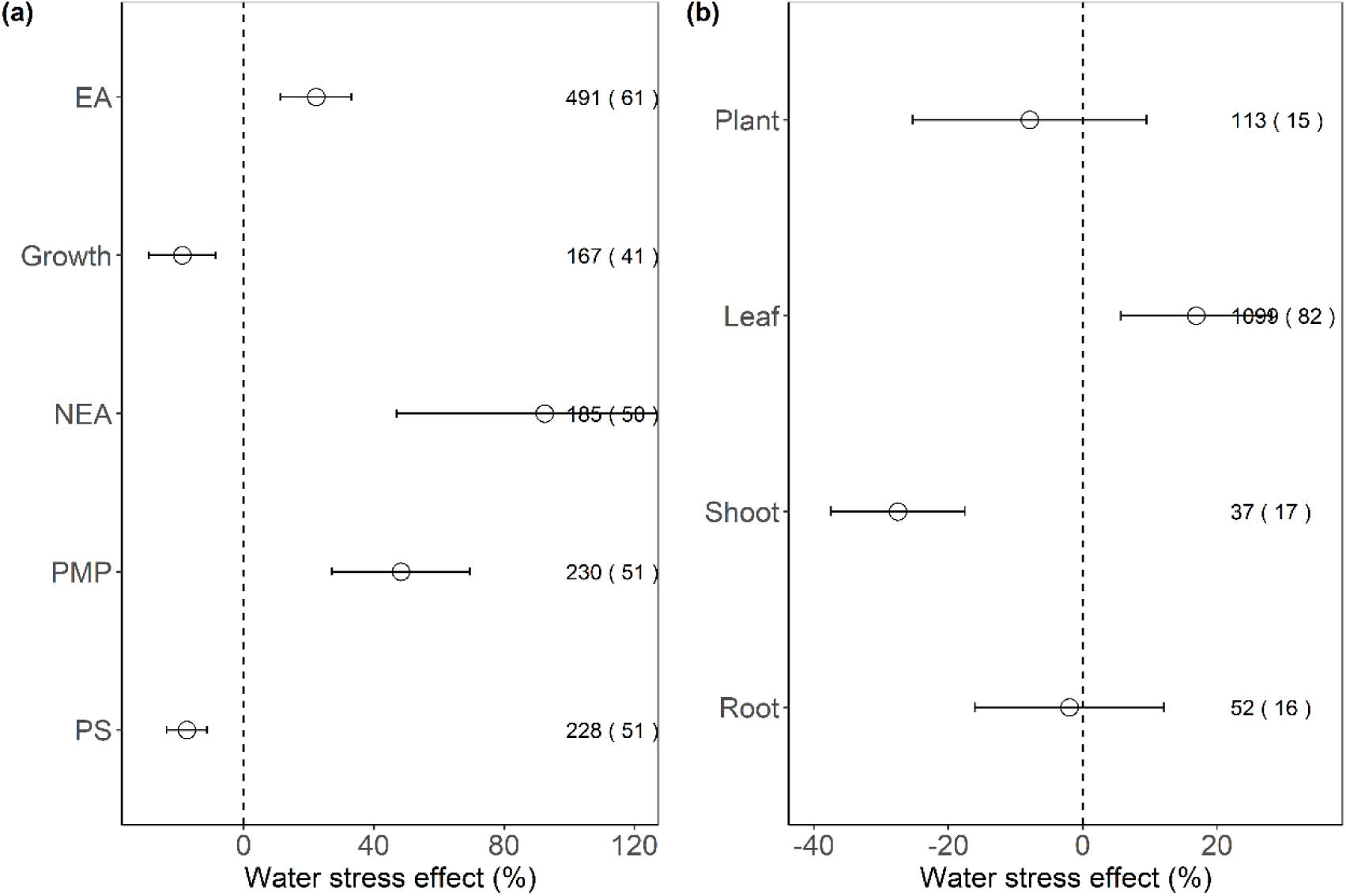
The response of plant performance (a) and tissues (b) to water stress. Values are means and 95% confidence intervals. Dashed black line is zero effect size. Numbers without and within parentheses represent the number of observations and studies, respectively. EA, NEA, PMP, and PS represent enzymatic antioxidants, non-enzymatic antioxidants, plasma membrane permeability, and photosynthesis, respectively.

With increasing water stress intensity, the effect size for ABA, CAT, EL, MDA, proline, and SOD increased significantly (all *P* < 0.05), whereas the effect size for Chl decreased (*P* < 0.01; Fig. 4). The effect sizes for APX, Chl, MDA, and SOD decreased significantly with experimental duration (all *P* < 0.05), and the effect size for AsA, EL, and protein increased (all *P* < 0.01; Fig. 5). The effect sizes for EA, NEA, and PMP increased significantly with water stress intensity (all *P* < 0.01; Fig. 6), and PMP decreased significantly with experimental duration (*P* < 0.001; Fig. 7).

**Figure 4.**
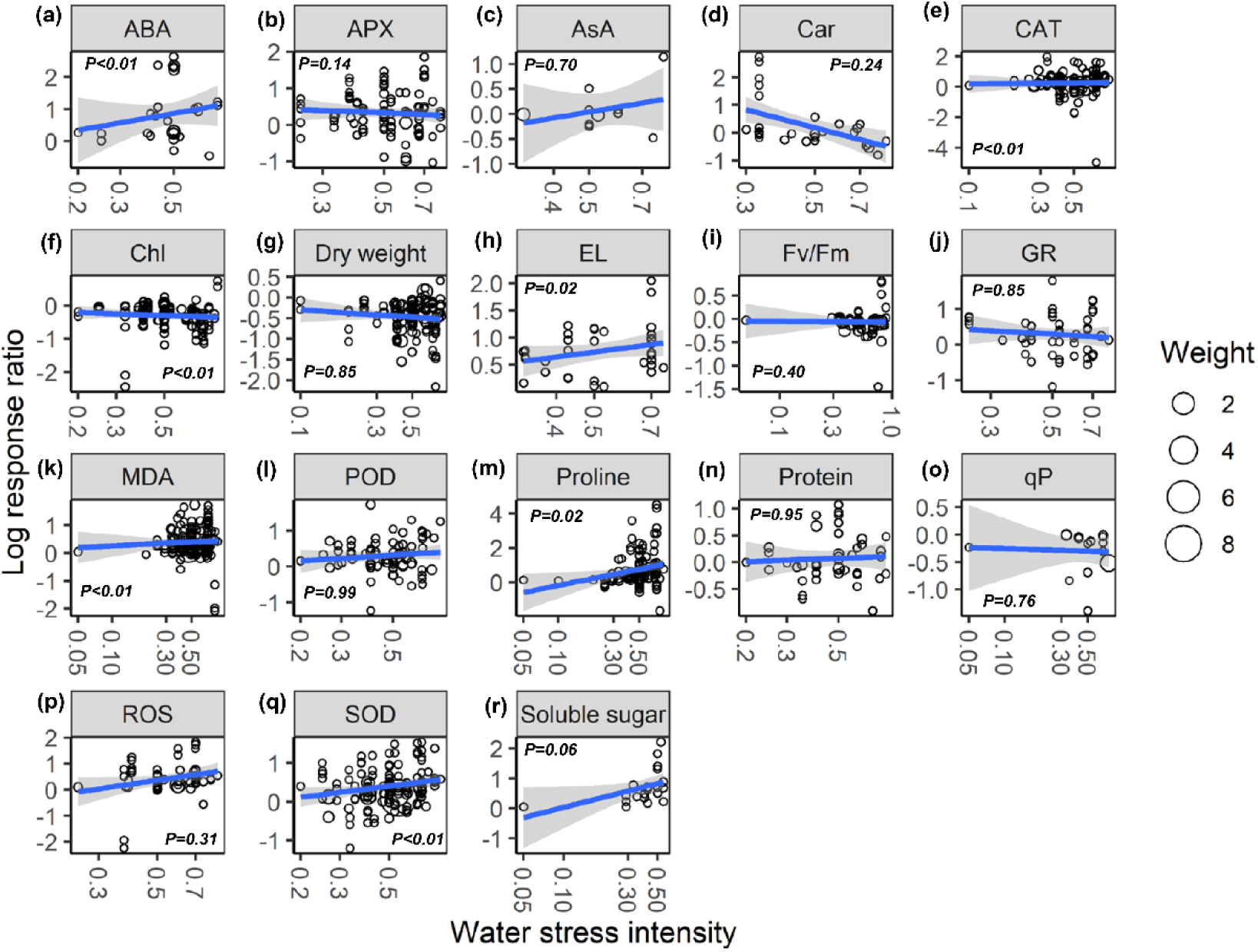
The responses of effect sizes of ABA (a), APX (b), AsA (c), Car (d), CAT (e), Chl (f), dry weight (g), EL (h), Fv/Fm (i), GR (g), MDA (k), POD (l), proline (m), protein (n), qP (o), ROS (p), SOD (q), and soluble sugar (r) to water stress intensity. Linear regressions are shown as solid blue lines, and 95% confidence intervals are the shaded areas. Circle sizes are proportional to the sampling variances. See Fig. 2 for abbreviations.

**Figure 5.**
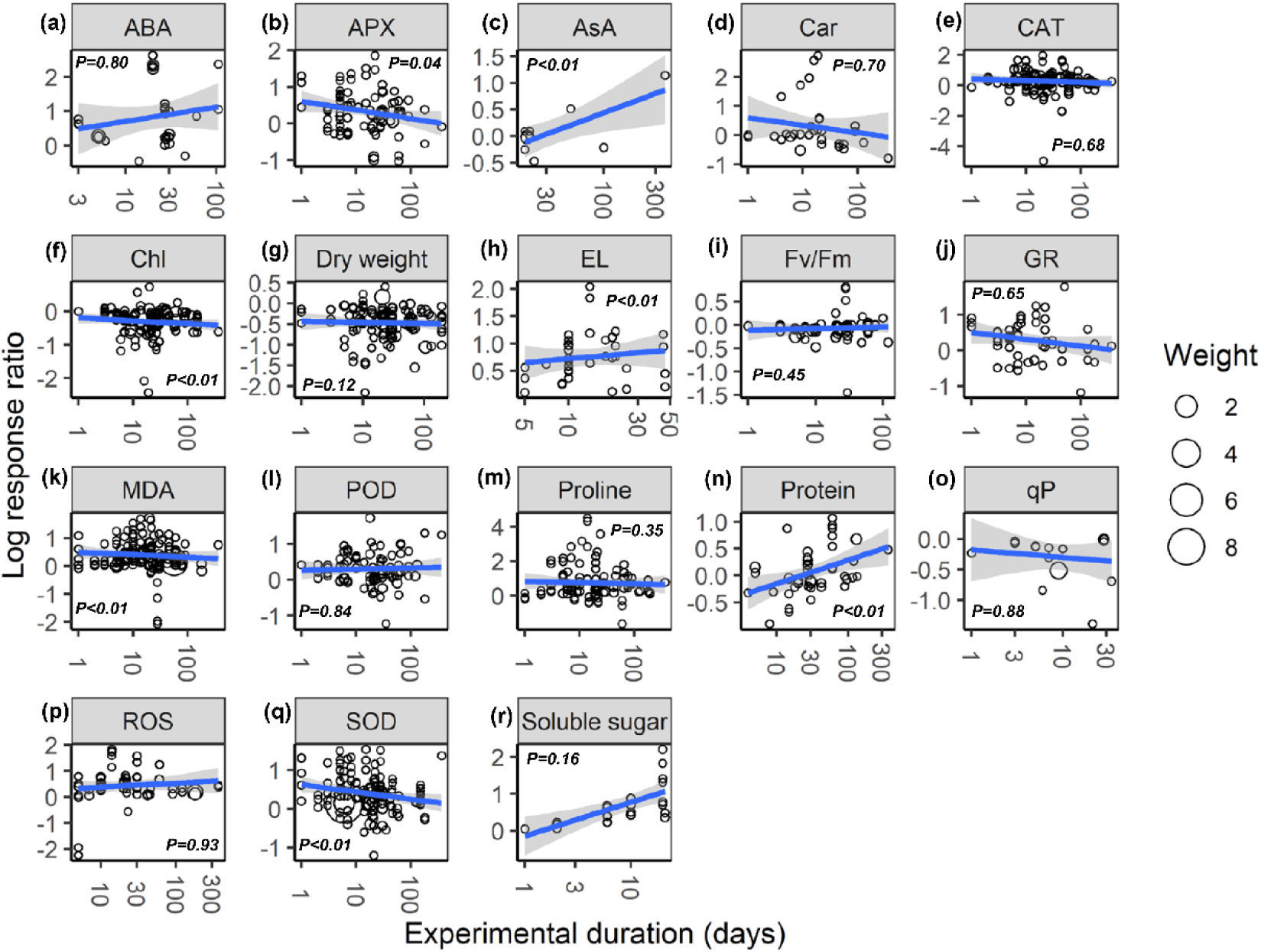
The responses of effect sizes of ABA (a), APX (b), AsA (c), Car (d), CAT (e), Chl (f), dry weight (g), EL (h), Fv/Fm (i), GR (g), MDA (k), POD (l), proline (m), protein (n), qP (o), ROS (p), SOD (q), and soluble sugar (r) to experimental duration. Linear regressions are shown as solid blue lines, and 95% confidence intervals are the shaded areas. Circle sizes are proportional to the sampling variances. See Fig. 2 for abbreviations.

**Figure 6.**
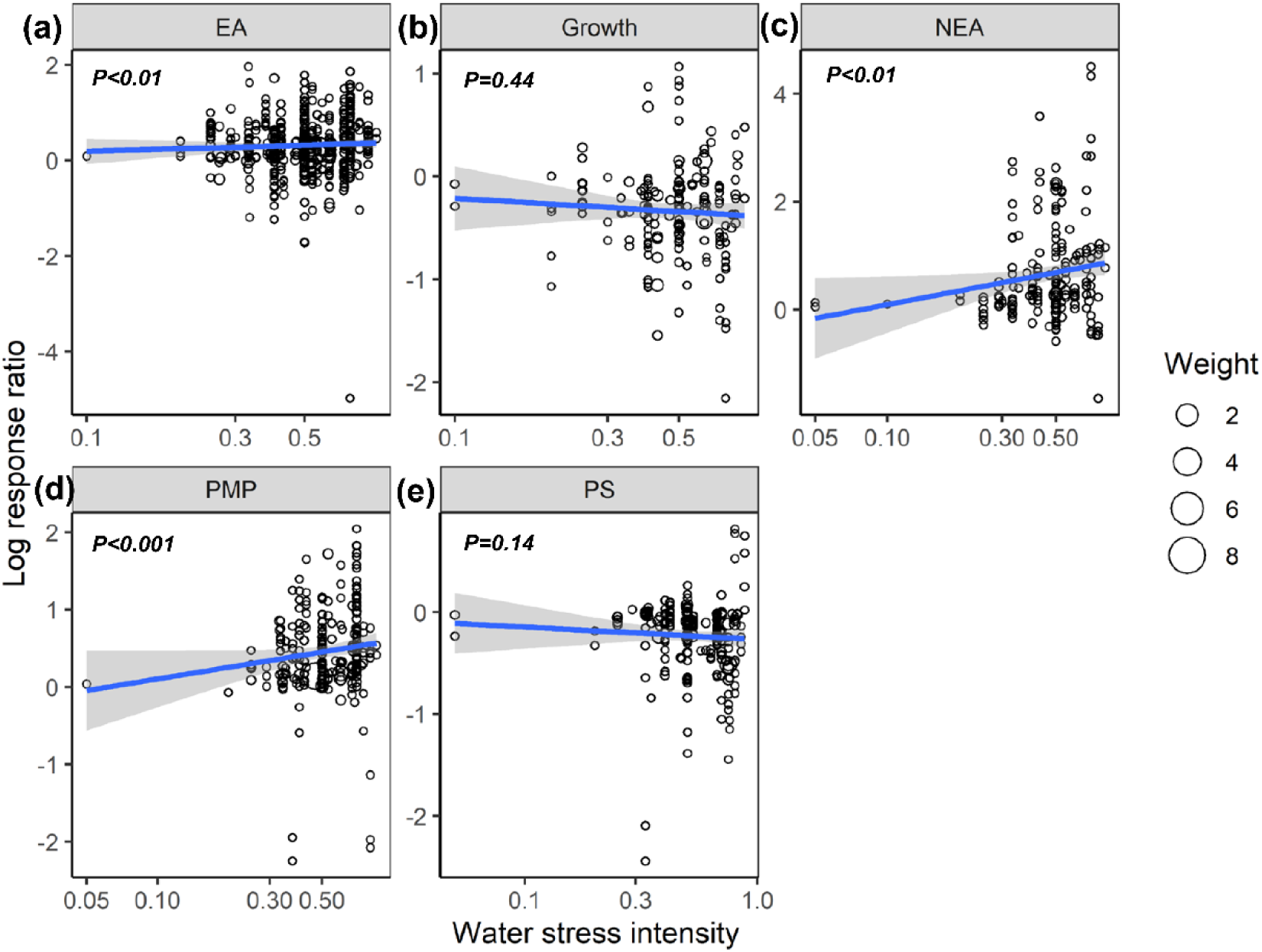
The responses of effect sizes of EA (a), growth (b), NEA (c), PMP (d), and PS (e) to water stress intensity. Linear regressions are shown as solid blue lines, and 95% confidence intervals are the shaded areas. Circle sizes are proportional to the sampling variances. EA, NEA, PMP, and PS represent enzymatic antioxidants, non-enzymatic antioxidants, plasma membrane permeability, and photosynthesis.

**Figure 7.**
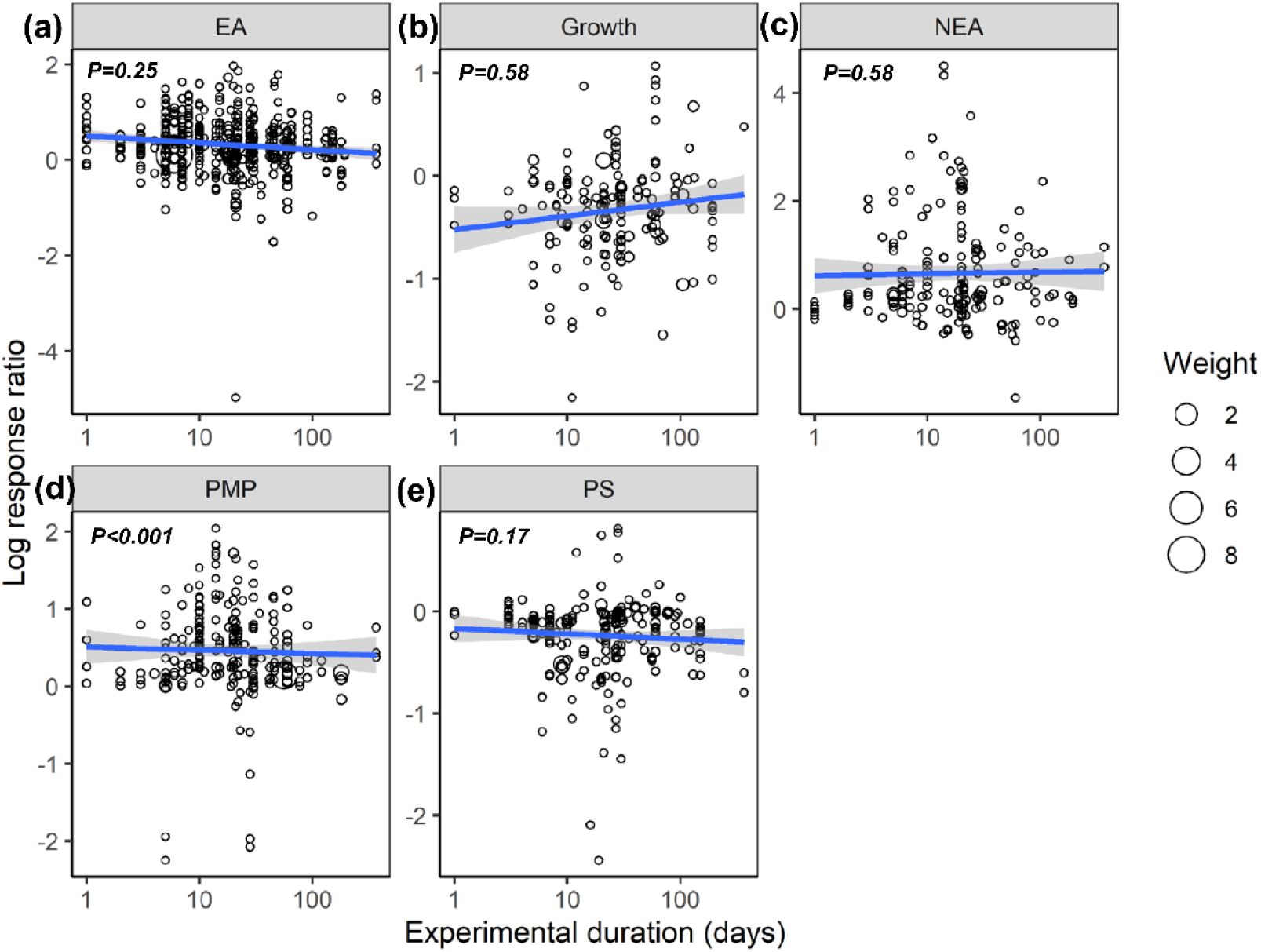
The responses of effect sizes of EA (a), growth (b), NEA (c), PMP (d), and PS (e) experimental duration. Linear regressions are shown as solid blue lines and 95% confidence intervals are the shaded areas. Circle sizes are proportional to the sampling variances. See Fig. 6 for abbreviations.

## 4. Discussion

The current meta-analysis based on 1301 observations, is the first to integrally examine the response patterns and mechanisms of plants to water stress on a global scale. A consistent and general response pattern of plants to water stress was found; these responses increased with water stress intensity but were mitigated with experimental duration. Below, we elaborate on the potential mechanisms for the observed patterns and conclude with suggestions for future research.

### 4.1. Mechanisms behind plant responses

As anticipated, our analysis revealed that water stress significantly inhibited plant growth and photosynthesis (Figs. 2, 3a). Further, we found that the negative response of Chl to water stress was more pronounced with increasing intensity and duration (Figs. 4f, 5f). This suggests that Chl is very sensitive to water stress among various types of plants. The probable explanation is that water stress can damage photosynthetic organs and alter leaf structures, thereby reducing the photosynthetic activities of plants and affecting plant growth (Aranjuelo et al., 2011).

The overproduction of ROS accompanied by increasing EL and MDA indicated a malfunction of the plasma membrane (Murray et al., 1989; Bouchemal et al., 2016) and lipid peroxidation (Sun et al., 2019), respectively. Our meta-analysis demonstrated that water stress significantly increased EL, MDA, ROS (Fig. 2), and stimulated PMP (Fig. 3a), which suggested that membrane damage occurred under water stress. Schneider et al. (2018) reported that high concentrations of ROS in plants are highly toxic to lipids and resulted in oxidative stress. Together, these results indicated that the overproduction of ROS was the primary mechanism of water stress. We also found that MDA and PMP exhibited positive responses to intensity, but negative responses to duration (Figs. 4k, 5k, 6d, 7d). One possible explanation is that plants have evolved some strategies to adapt to water stress (see the discussion below) (Anjum et al., 2012; Sperdouli and Moustakas, 2012). While increases in the EL of plant cells to both intensity and duration were observed (Figs. 4h, 5h), this index should be useful in screening for drought-resistant species.

Our study offers new insights into the increase of EA being associated with scavenged ROS under water stress (Figs. 2, 3a, 6a). Although CAT, POD, and SOD activities were higher under water stress than the control, both APX and GR did not show significant responses to water stress. This suggested that the EA associated with the Halliwell-Asada pathway may work less efficiently than CAT, POD, and SOD, likely because various enzymes located within different cellular compartments have disparate functions (Mittler, 2002; Gill and Tuteja, 2010; Sun et al., 2019). Further, the CAT and SOD increased with intensity (Figs, 4e, q), being attributed to the enhanced tolerance to stress (Caverzan et al., 2016), which maintained normal metabolic processes (Schneider et al., 2018). Thus, we recommend that the kinetics involved in the enzymatic responses to water stress should be investigated in future experiments.

Our study also revealed higher ABA, AsA, proline, soluble sugar, and NEA (Figs. 2, 3a) under water stress. We also found that the positive responses of ABA, proline, and NEA were more pronounced with intensity (Figs. 4, 6c), which suggested that ABA and proline were sensitive plant parameters to water stress. Higher concentrations of ABA acted to adapt to water stress (Belkheiri and Mulas, 2013), where increased proline accumulation was considered to mitigate the adverse effects of ROS (Chen and Dickman, 2005). Interestingly, plant roots were not significantly impacted by water stress (Fig. 3b). One possible explanation is that increases in the numbers of root ducts improved the efficacy of water transport; thus assisting plants to resist water stress (Lee et al., 2016).

### 4.2. Suggestions for future experiments

We encountered two important inconsistencies in our meta-analysis across studies. Firstly, only 10 studies of the 84 in our dataset lasted more than three months. Experimental duration is a vital factor that affects plant responses to water stress (Figs. 5, 7); therefore, we suggest that longer time scales (> three months) be employed in future experiments. Secondly, as with several ecological meta-analyses, we found a hemispheric bias in our knowledge of the effects of water stress on plants (Feeley et al., 2017). Most observations were derived from experiments performed at a latitude of > 19.3° in the Northern Hemisphere (only nine studies were conducted in the Southern Hemisphere). Therefore, we are hopeful that our study will motivate new publications in underrepresented regions (Fig. 1).

## 5. Conclusion

In conclusion, our meta-analysis, which employed a global empirical dataset for the responses of plants to water stress, revised the previous notion that water stress inhibits plant growth (Schneider et al., 2018). Divergent responses of Chl, EL, and ROS were partly explained by the inhibition of plant growth. Water stress affected plant performance primarily through the overproduction of ROS, which led to plasma membrane damage. Meanwhile, a variety of physiological indexes, i.e., CAT, SOD, ABA, and proline were activated to control the levels of cellular ROS to compensate for water stress. These above indexes were evaluated to facilitate screening for drought-resistant species. Further, the effects of water stress were observed to be more pronounced with intensity, which was, however, mitigated with duration. Therefore, imbuing plants with the capacity to scavenge excessive ROS will be useful in the future to facilitate their endurance during drought events.

## Abbreviations

ABA: abscisic acid;
APX: ascorbate peroxidase;
AsA: ascorbate;
Car: carotenoid;
CAT: catalase;
Chl: chlorophyll;
EA: enzymatic antioxidants;
EL: electrolyte leakage;
Fv/Fm: maximal efficiency of PSII photochemistry;
GR: glutathione reductase;
MDA: malondialdehyde;
NEA: non-enzymatic antioxidants;
PMP: plasma membrane permeability;
POD: peroxidase;
PS: photosynthesis;
qP: photochemical quenching coefficient;
ROS: reactive oxygen species;
SD: standard deviation;
SOD: superoxide dismutase.

## Appendix A. Supplementary data

Additional supporting information that accompanies this paper may be found in the online version of this article.

All data and R codes will be published on figshare once the manuscript is accepted for publication.

